# Syndiniales parasites drive species networks and are a biomarker for carbon export in the oligotrophic ocean

**DOI:** 10.1101/2023.06.29.547083

**Authors:** Sean R. Anderson, Leocadio Blanco-Bercial, Craig A. Carlson, Elizabeth L. Harvey

**Affiliations:** Department of Biological Sciences, University of New Hampshire, Durham, NH, USA; Marine Chemistry and Geochemistry Department, Woods Hole Oceanographic Institution, Falmouth, MA, USA; Bermuda Institute of Ocean Sciences, Arizona State University, St. George’s, Bermuda; Department of Ecology, Evolution and Marine Biology and the Marine Science Institute, University of California, Santa Barbara, CA, USA

## Abstract

Microbial associations that result in phytoplankton mortality are important for carbon transport in the ocean. This includes parasitism, which in microbial food webs, is dominated by the marine alveolate group, Syndiniales. Parasites are expected to contribute to carbon recycling via host lysis; however, knowledge on host dynamics and correlation to carbon export remain unclear and limit the inclusion of parasitism in biogeochemical models. We analyzed a 4-year 18S rRNA metabarcoding dataset (2016-2019), performing network analysis for twelve discrete depths (1- 1000 m) to determine Syndiniales-host associations in the seasonally oligotrophic Sargasso Sea. Analogous water column and sediment trap data were included to define environmental drivers of Syndiniales and their correlation with particulate carbon flux (150 m). Syndiniales accounted for 48-74% of network edges, most often associated with Dinophyceae and Arthropoda (mainly copepods) at the surface and Rhizaria (Polycystinea, Acantharea, and RAD-B) in the aphotic zone. Unlike other major groups, Syndiniales were significantly (and negatively) correlated with particulate carbon flux, suggesting parasites may drive flux attenuation through remineralization. Examination of Syndiniales amplicons revealed a range of depth patterns, including specific ecological niches and vertical connection among a subset (19%) of the community, the latter implying sinking of parasites (infected hosts or spores) on particles. Our findings point to the use of Syndiniales as biomarkers of carbon export, highlighting their importance for marine food webs and biogeochemistry.

**Significance Statement:** Syndiniales parasites are widespread in the ocean and represent a potentially important, albeit poorly resolved, source of carbon recycling. Here, we assess Syndiniales population dynamics, trophic relationships, and links to carbon export in the Sargasso Sea. Species networks at all depths were driven by Syndiniales, with parasite-host relationships varying with depth based on shifts in host composition. Syndiniales were the only eukaryote group to be significantly (and negatively) correlated with particulate carbon flux, indicating their contribution to flux attenuation via remineralization. Yet, a subset of parasites was vertically connected between photic and aphotic zones, suggesting continued export. Our findings elevate the critical role of Syndiniales in marine microbial systems and reveal their potential use as biomarkers for carbon export.

## Introduction

Phytoplankton are central to the biological carbon pump, fixing atmospheric carbon dioxide and contributing to the sinking of organic matter (1). The biological carbon pump exports ∼10 Pg C y^-1^ from the surface oceans, with contributions from many biological, chemical, and physical processes (2, 3). Particulate organic carbon (POC) fixed by primary producers is exported via several well-known pathways: gravitational sinking, aggregation, active transport from diel migrators, and physical mixing (4–6). Phytoplankton mortality, and the species interactions that underpin it, also drive export strength and efficiency (7, 8). However, mechanistic links between species interactions and carbon export remain elusive and preclude incorporation of mortality in ocean carbon export models (9, 10). Insight into microbial interactions is necessary to define the contribution of plankton mortality to carbon cycling and accurately resolve the ecological mechanisms that drive carbon export.

Parasitism is arguably the most common lifestyle employed by organisms on Earth (11), and yet in the ocean, it is rarely incorporated in food web or biogeochemical models (12). Routine metabarcoding surveys have revealed the hidden diversity and global distribution of protist parasites in the ocean (13, 14). Marine alveolates in the group Syndiniales are the most ubiquitous and phylogenetically diverse, with sequences observed in virtually all marine biomes (15–17). In addition, several recent studies have reported a high proportion of Syndiniales reads in particles collected from sediment traps (10, 18–21). Yet, direct links between Syndiniales and carbon export have not been established, making it difficult to contextualize the role of parasites in carbon cycling.

Carbon exported to the deep ocean is partly mediated by organismal interactions (7, 22), which for parasites, have not been well defined vertically in the water column (23). Syndiniales infect a wide range of hosts (e.g., dinoflagellates, ciliates, radiolarians, and copepods), with several studies reporting top-down pressure on coastal phytoplankton blooms that rival grazing loss (15, 24). Infection begins with attachment of a motile parasite spore (<10 µm) to a host cell, followed by rapid digestion of host material and growth inside the nucleus (25–27). After 2-3 days, whereby the parasite increases in volume 200-fold, the host ruptures, and releases hundreds of new spores into the environment (27). It is estimated that up to 70% of host biomass is released as labile dissolved organic matter (DOM) that can be recycled by heterotrophic bacteria (28). Estimates of parasite-released DOM, and its lability, likely depend on host community composition and biomass (29), as well as environmental conditions, like nutrient concentrations and temperature (30, 31). Nevertheless, building evidence suggests Syndiniales are central to microbial food webs and carbon cycling, making it imperative to detail host associations with depth and their involvement in carbon export.

Here, we investigate Syndiniales infection dynamics throughout the photic (1-120 m) and aphotic (160-1000 m) zones at the Bermuda Atlantic Time-series Study (BATS) site, a long-term (>30 y) ocean monitoring program in the seasonally oligotrophic Sargasso Sea (32). Recent 18S metabarcoding work at BATS revealed a high relative abundance of Syndiniales (∼40%) throughout the water column (33) and their presence in sediment traps (20). We performed covariance network analysis of a 4-year (2016–2019) 18S rRNA metabarcoding dataset (33), constructing networks for each of twelve discrete depths (1-1000 m). Our analysis revealed depth-dependent trends in Syndiniales networks and putative relationships. Sediment trap data collected from the same location revealed a significant and negative correlation between Syndiniales and bulk POC flux at 150 m, implying increased parasite relative abundance may enhance flux attenuation through remineralization of host carbon. In addition, we found evidence of vertical transport among a subset (19%) of the parasite community, matching the percent of primary production (10-20%) exported from the surface oceans each year.

## Results and Discussion

### Spatiotemporal variability and environmental drivers of Syndiniales

A total of 18,643 unique Syndiniales amplicon sequence variants (ASVs) were identified over the entire 18S dataset, which included monthly samples collected over 4 years (2016–2019) and twelve discrete depths (1-1000 m). Syndiniales were consistently distributed with depth and time (33), accounting for 26-59% of total 18S reads (Fig. 1A; SI Appendix, Fig. 1). Other studies in oligotrophic systems have found Syndiniales comprise a significant proportion of the eukaryotic population in deep waters (17, 34, 35) and sediment traps (18, 36). By comparison, relative abundance of other major 18S groups were more variable with depth (33), with higher Dinophyceae and Arthropoda (mainly copepods) relative abundance in the photic zone that shifted to a Rhizaria-dominated community in the aphotic zone and included several radiolarians like Polycystinea, Acantharea, and RAD-B (Fig. 1A; SI Appendix, Fig. 1).

**Figure 1.**
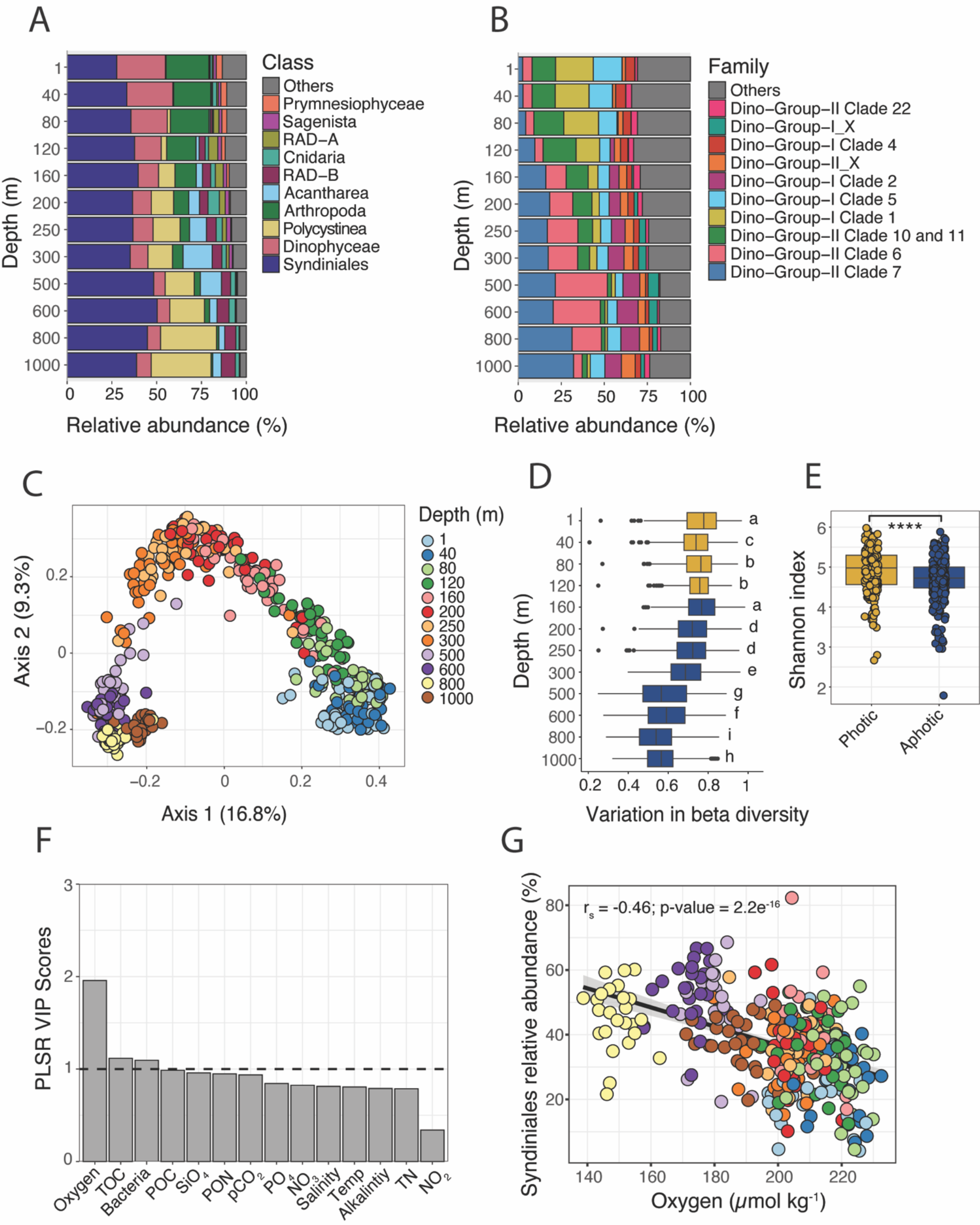
Syndiniales population dynamics and environmental drivers at BATS. Stacked bar plots of 18S relative abundance at the class level (**A**) and within Syndiniales (**B**) at the family (clade) level at each depth. Stacked bar plots display the top ten groups based on relative abundance (“Others” in gray). Values are presented as the mean at each depth; n = 43-53 (C) Principal coordinates analysis based on Bray-Curtis dissimilarity of all samples, filtered to only include Syndiniales ASVs. Samples represent single replicates and are colored by depth. The proportion of variance explained by the first two axes is indicated on the plot. (**D**) Variation in beta diversity as a function of sampling depth. Values represent the mean ± SD (n = 43-53), with letters indicating statistical differences. Samples are colored by their position in the photic (gold) vs. aphotic zones (blue). (**E**) Shannon index values for Syndiniales ASVs in the photic (n = 206) vs. aphotic zones (n = 381). A Wilcoxon test was used to estimate significant differences in the mean (**** = p-value < 0.0001). (**F**) Variable influence on the projection (VIP) scores for each factor estimated from the partial least squares regression (PLSR) model and ordered from highest to lowest. VIP > 1 (dashed line) indicate variables most important to the model. Oxygen (µmol kg^-1^); TOC = total organic carbon (µmol kg^-1^); Bacteria = bacteria cell density (10^8^ cells kg^-1^); POC = particulate organic carbon (µg kg^-1^); SiO4 = silicate (µmol kg^-1^); PON = particulate organic nitrogen (µg kg^-1^); *p*CO2 = partial pressure of carbon dioxide (µatm); PO4 = phosphate (µmol kg^-1^); NO3 = nitrate (µmol kg^-1^); Salinity (psu); Temp = temperature (°C); Alkalinity (µmol kg^-1^); TN = total nitrogen (µmol kg^-1^); NO2 = nitrite (µmol kg^-1^). (**G**) Spearman rank correlation (rs) between Syndiniales relative abundance vs. oxygen concentration. All samples were considered and colored by depth. Regression line and 95% confidence intervals are shown.

Syndiniales were largely assigned to Dino-Groups I and II (SI Appendix, Fig. 2), which are considered the two most prevalent and diverse Syndiniales groups in the ocean (15). Depth variability was most apparent at the family (clade) level (Fig. 1B). Several clades, like Dino-Group-II Clade 10+11 and Dino-Group-I Clades 1 and 5, had higher relative abundance in the photic zone, while Dino-Group-II Clades 6 and 7 and Dino-Group-I Clade 2 were more prevalent in the aphotic zone (Fig. 1B). Clade-level patterns observed at BATS align closely with previous observations of Syndiniales distribution in the water column (15, 36), further supporting the presence of depth-dependent ecological niches within Syndiniales. As is the case with metabarcoding data, the relative abundance of many groups, including Syndiniales, may be inflated due to high 18S gene copy number (37). We focused our analysis on depth-related trends within Syndiniales and note that other groups at the surface (e.g., dinoflagellates) were also biased by potential copy number.

**Figure 2.**
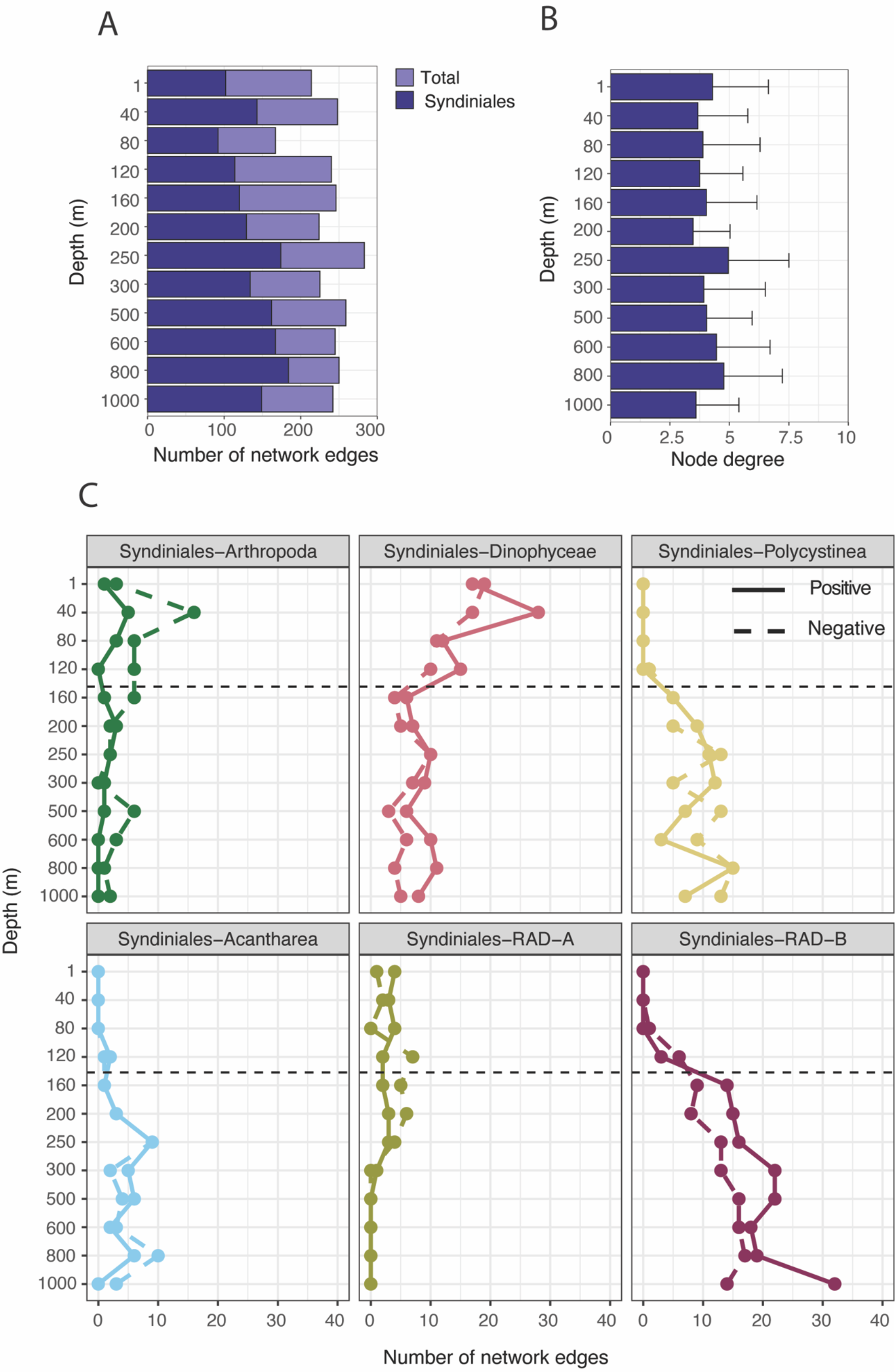
Results from SPIEC-EASI network analysis that considered the top 150 18S amplicon sequence variants (ASVs) at each depth. (**A**) The number of network edges (associations) for the total networks (light purple) and those attributed to Syndiniales ASVs (dark purple). (**B**) Syndiniales node degree for each depth network, i.e., the number of other 18S ASVs connected to a single Syndiniales node (or ASV). Values represent the mean ± SD at each depth. (**C**) Depth-related trends display the number of positive (solid) and negative (dashed) edges between Syndiniales and putative host groups at the class level (Arthropoda, Dinophyceae, Polycystinea, Acantharea, and RAD-A/B). The transition from photic to aphotic zones are distinguished with dashed lines at 140 m (0.1% light). Network edges filtered for Syndiniales are in Dataset S2.

Depth, and not time, was important in structuring Syndiniales community composition at BATS (Fig. 1C-E; SI Appendix, Fig. 3). Other 18S metabarcoding studies at this site (33, 38), and in the North Pacific (17), have noted enhanced depth effects over seasonal cycles in driving protist communities. Syndiniales were significantly clustered by depth (PERMANOVA R = 0.13; p- value < 0.001), with communities being more tightly clustered and less diverse in the aphotic zone (Fig. 1C-E). Apart from seasonal variability at 1 m (PERMANOVA R = 0.2; p-value < 0.001), Syndiniales were not significantly influenced by seasonal effects (SI Appendix, Fig. 3). Seasonality in the Sargasso Sea is characterized by deep convective mixing (∼150-300 m) between January-March and stratification of the water column (mixed layer depth <20 m) as early as May (32), driving seasonal patterns in phytoplankton, heterotrophic bacteria, and viruses (39–41). Enhanced winter mixing typically facilitates higher plankton biomass, net primary production, and carbon export to 150-300 m (32, 42). Seasonal effects on Syndiniales may be minimized in the mixed layer due to their ubiquity, host range, and tolerance for variable physical and chemical conditions. Others have noted seasonality among Syndiniales in coastal time series with weekly resolution (43, 44). Sampling intervals were less resolved (monthly) at BATS, which may have masked temporal effects.

**Figure 3.**
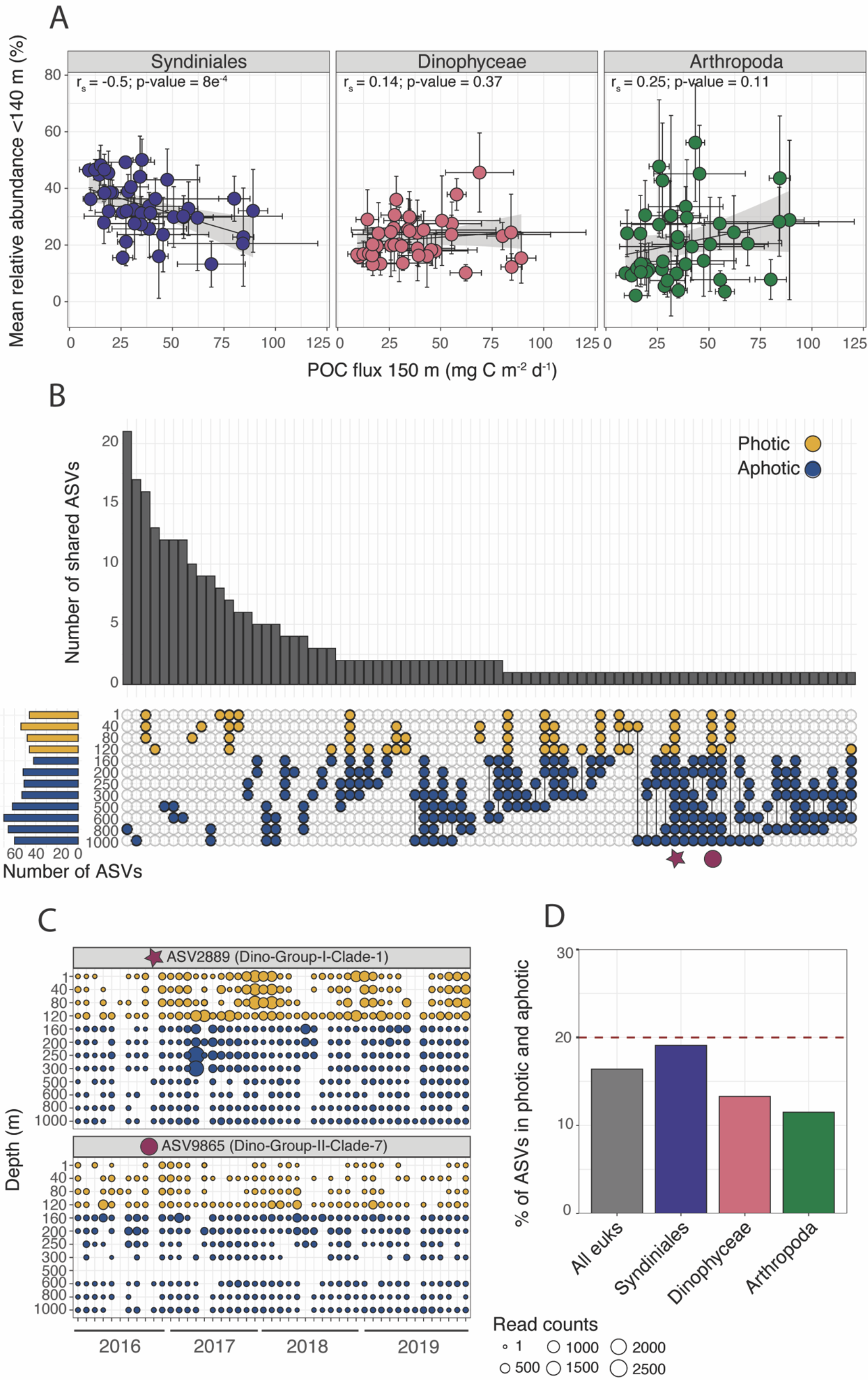
Syndiniales and POC flux from 150 m at BATS. (**A**) Spearman rank correlations (rs) between Syndiniales, Arthropoda, or Dinophyceae mean relative abundance in the photic zone (< 140 m) vs. mean bulk POC flux estimated from sediment traps deployed at 150 m. Regression line and 95% confidence intervals are shown. (**B**) UpSet plot to visualize the intersection of Syndiniales ASVs between depth networks. Horizontal bars on the left show the number of Syndiniales ASVs within each depth network. Points and lines between points indicate the intersection, while bar plots on the top panel represent the number of shared Syndiniales ASVs. Networks are colored by their position in the photic (gold) vs. aphotic zones (blue) to better display vertical connection. Overlap is based on presence-absence data. (**C**) Example profiles of specific Syndiniales ASVs from the intersection plot (red star and circle) to show spatial and temporal changes in rarefied reads counts at the ASV level. (**D**) The percentage of ASVs for all eukaryotes, Syndiniales, Dinophyceae, and Arthropoda present at any time in both the photic (1-120 m) and aphotic (160-1000 m) zones. The red dashed line indicates upper limit estimates of percent of primary production exported from photic zone each year (66).

Partial least squares regression (PLSR) was used to identify the most important environmental variables that influenced Syndiniales read counts at BATS (Fig. 1F). Vertical profiles of hydrographic data, nutrient concentrations, and carbon measurements (SI Appendix, Fig. 4; Dataset S1) were typical for this site (42). Oxygen concentration was most important in explaining Syndiniales read counts (Fig. 1F) based on variable importance on the projection (VIP) scores from the PLSR model (SI Appendix, Fig. 5). Other variables were also important (VIP > 1), including total organic carbon (TOC) concentration and bacterioplankton cell density (Fig. 1F). Syndiniales relative abundance was negatively correlated with oxygen concentration (Spearman rs = -0.46; p-value < 0.001), with highest abundance in the aphotic zone where oxygen was <180 µmol kg^-1^ (Fig. 1G). Syndiniales are known to persist and contribute to species interactions in low oxygen environments (23) and other extreme marine habitats, like hydrothermal vents (45).

### Depth-specific networks and putative parasite-host relationships

Co-occurrence networks are often applied to amplicon sequencing data to infer species relationships in the ocean (31, 46). Network analysis is a powerful, hypothesis-driven approach, particularly as many marine microbial species remain uncultured and have unrealized ecological functions (47, 48). Caution should be used when interpreting network results, as correlations cannot be defined as true ecological interactions (49). Networks at BATS, which included the top 150 most abundant 18S ASVs from each depth, were dominated by Syndiniales (Fig. 2A). A total of 268 Syndiniales ASVs accounted for 48-74% of 18S edges across networks, with a slight increase in edge number with increasing depth (Fig. 2A). Other sequence-based studies have noted a large contribution of Syndiniales ASVs to species edges inferred by co-occurrence networks (46, 50, 51). Within networks, a single Syndiniales ASV was often connected to more than two putative host ASVs on average, with little variation with depth (Fig. 2B). Similar patterns were observed for potential host ASVs connected to different parasites (SI Appendix, Fig. 6), with node degree often > 2. However, node degree of potential host groups was vertically structured at times, especially for Arthropoda, with a peak in node degree at 120-200 m (SI Appendix, Fig. 6). These findings imply parasite-host dynamics at BATS may lean towards opportunistic strategies that are depth-dependent, though additional quantitative work is needed to confirm infection patterns and host specificity under different conditions (48).

Syndiniales were most often associated with several eukaryotic groups, like Arthropoda, Dinophyceae, and Rhizaria (Polycystinea, Acantharea, and RAD-A/B), all representing verified or potential hosts of the parasite (15). The number of network edges mirrored depth-specific trends in relative abundance among putative hosts (Fig. 1A; Fig. 2C). For example, the number of edges between Syndiniales-Arthropoda and Syndiniales-Dinophyceae were highest in the photic zone, while associations with Polycystinea, Acantharea, and RAD-B were elevated in the aphotic zone (Fig. 2C). The exception were associations between Syndiniales and RAD-B that were most prevalent among radiolarians in deeper waters, despite RAD-B accounting for <10% of sequence reads (Fig. 1A; Fig 2C). Positive and negative edges for each respective pairing were tightly linked with depth (Fig. 2C), with some exceptions (e.g., increase in negative Syndiniales-Arthropoda edges at 40-160 m). Interpreting the sign of network edges is difficult without further experimental context (52). Positive edges may indicate infection (copresence) while negative edges may represent lysis that separates groups (mutual exclusion); however, edges can also imply an overlapping or different ecological niche.

Dinoflagellates are the most well-studied hosts of Syndiniales (53), with infection reported in the lab and field, including in genera revealed here, like *Gyrodinium*, *Gymnodinium*, and *Prorocentrum* (SI Appendix, Fig. 7; Dataset S2). Interactions between Syndiniales and other hosts, like copepods and radiolarians, are less clear. Copepods, which in our networks mostly involved *Calocalanus, Clausocalanus, and Triconia* (SI Appendix, Fig. 7; Dataset S2), may be directly infected by Syndiniales Groups I and IV (54), though it is more likely copepods are associated with parasites via uptake of infected prey (55). Direct infection has not yet been observed in radiolarians. However, many have speculated on a parasitic relationship (15, 36, 56), given their shared ecological niche in deep waters and the presence of Syndiniales sequences in single-cell radiolarian isolates (57, 58). Our network analysis supports the role of radiolarians as potentially important hosts of Syndiniales in the aphotic zone. We identified several putative parasite-host relationships among radiolarians (SI Appendix, Fig. 7; Dataset S2), involving Group I (Clade 2) and II (Clades 6 and 7) parasites correlated to Polycystinea (*Cladococcus* and *Heliosphaera*), Acantharea (Acantharea Group I, *Acanthoplegma*, and *Litholophus*), and RAD-B (Groups Ia, Ib, II, and III). Given the global abundance of radiolarians and their contribution to carbon export (59), it will be critical to further explore parasitic infection in this group.

### Role of Syndiniales in POC flux and vertical transport

Sediment trap data collected at BATS was included in our analysis to explore links between POC flux (at 150 m) and integrated Syndiniales relative abundance in the photic zone (1-140 m). Bulk POC flux at 150 m ranged from 9.4 to 89.1 mg C m^-2^ d^-1^ over the 4-year dataset. During this time, minimal seasonality was observed, except for significantly lower mean POC flux in the fall (22.6 mg C m^-2^ d^-1^) compared to other seasons (40.5-46 mg C m^-2^ d^-1^). Previous work has reported higher POC flux in spring vs. fall at sediment traps at 150, 200, and 300 m, resulting from increased phytodetrital and fecal matter aggregation in response to elevated primary production (20, 32). We observed a significant negative correlation (Spearman rs = -0.51; p-value = 0.001) between bulk POC flux at 150 m and mean Syndiniales relative abundance in the photic zone (Fig. 3A), a trend that was not significant for other groups with high relative abundance (Dinophyceae or Arthropoda). In addition to POC flux at 150 m, Syndiniales relative abundance was significantly (and negatively) correlated with TOC and POC concentration (Spearman rs = - 0.4 and -0.32; p-values < 0.001). Parasites are thought to have a similar biogeochemical impact as viruses, rerouting carbon away from POC and into pools of labile dissolved and particulate organic matter (DOM/POM) that can fuel bacterial production (28). Our findings support the expected ecological role of protist parasites in the ocean (26), and suggest that Syndiniales in particular, significantly drive particle flux attenuation through remineralization of host carbon.

Yet, the contribution of parasitism to DOM remains unclear in microbial systems (60) and warrants deeper investigation through ‘omics-based surveys and controlled lab experiments.

Our results also suggest that a proportion of Syndiniales are exported out of the surface, based on the presence of Syndiniales ASVs in both the photic and aphotic zones (Fig. 3B-D). Indeed, Syndiniales ASVs are consistently reported within sediment traps across diverse marine systems (18, 19, 36). While our work cannot conclude the exact mechanisms of export, Syndiniales may be contained within sinking phytodetritus, fecal aggregates, or fecal pellets (10), all of which constitute the bulk of sinking POC at BATS (20). Infected hosts may be transported to deep waters via zooplankton fecal matter, expedited through diel migration (61). Though infection is rapid (2-3 d), motile parasite spores can survive outside the host for up to 15 d under laboratory conditions (27). Therefore, it may be possible for spores to survive on sinking aggregates, which at BATS range in size from 60-1862 µm (20, 62). There is also evidence that Syndiniales become more abundant on particles over time, indicating active infection of attached hosts (10). Spores may survive without their preferred hosts, by switching hosts, producing cysts (63), or acquiring alternative energy sources. Similar adaptive strategies have been observed among prey-limited heterotrophic dinoflagellates (64). Lastly, Syndiniales may reach deep waters through convective mixing (42, 65), either directly or on particles.

Though negatively linked with POC flux at 150 m, 19% of Syndiniales ASVs showed signs of vertical transport at BATS (Fig. 3B-D). Compared to Syndiniales, fewer ASVs (12-16%) were present throughout the water column for Arthropoda, Dinophyceae, and all eukaryotes (Fig. 3D), suggesting less vertical connection. Moreover, Syndiniales ASVs in the networks were present across seasons (SI Appendix, Fig. 8), with signals of vertical transport consistent over the 4-year period and seasonally recurrent (Fig. 3C). As demonstrated here and hinted by others (15, 26), Syndiniales parasites likely play an important role in modulating sinking POC and may serve as biomarkers for carbon export. As an example, the percent of sinking parasite ASVs at BATS is concurrent with the percent of primary production (10-20%) sinking each year from the photic zone (66). While not directly comparable and requiring additional evidence, our results suggest the potential application of molecular tools to capture export by evaluating Syndiniales ASVs and their correlation to analogous flux estimates. Given the dominance of Syndiniales relative abundance in the surface ocean, at depth, and in sediment traps, we propose using Syndiniales parasites as a biomarker for carbon export flux in the ocean. Additional field and laboratory work should focus on characterizing sinking mechanisms, distribution of infected hosts and spores in the water column and on particles (e.g., sample fractionation; (44)), and changes in parasite-host dynamics with respect to biogeochemistry over space and time.

## Conclusion

The influence of parasitism on marine microbial food webs and carbon cycling is unclear. This study provides detailed information on depth-specific population dynamics and trophic relationships among the widespread protist parasite, Syndiniales, and links parasites with POC flux and vertical transport of sinking POC in the oligotrophic North Atlantic Ocean. Syndiniales were connected to a range of known and putative hosts, like dinoflagellates, copepods, and radiolarians, with associations varying with depth based on changes in host composition. A strong and negative correlation between Syndiniales in the photic zone and POC flux to 150 m implies parasites are contributing to POC flux attenuation and carbon recycling as particles move from the photic to aphotic zones, an ecological role that has been intimated by others (26).

Syndiniales communities in the aphotic zone may represent resident clades that are adapted to specific ecological niches (15), as well as a subset of parasites that sink and may exhibit opportunistic infection strategies. Given the strong connection with POC flux from the photic zone, it may also be expected that parasites represent important sources of DOM at depth that can fuel microbial communities (45). In total, these findings elevate the role of Syndiniales in marine food webs, highlighting their importance to species networks at all depths and potential use as biomarkers for carbon export.

## Materials and Methods

### Sample collection

Monthly seawater samples were collected at BATS over a 4-year period (February 2016 to December 2019) at twelve discrete depths in the water column (1, 40, 80, 120, 160, 200, 250, 300, 500, 600, 800, and 1000 m). At each depth and month, 4 L of seawater was filtered through 0.22 µm Sterivex filters (MERCK, MA, USA) and filters were stored at -80°C (33). Only a single replicate filter was collected. Each profile was retrospectively assigned to one of four seasons (33) that corresponded to the position of the mixed layer depth (e.g., mixed in winter vs. stratified in summer). CTD profiles (temperature, oxygen, and salinity) and discrete chemical and biological data were also collected each month and considered here to relate to Syndiniales relative abundance. Dissolved nutrients (NO3, NO2, PO4, SiO4), total organic carbon (TOC), total *p*CO2, particulate organic carbon and nitrogen (POC and PON), total alkalinity, total nitrogen, and bacteria cell density were included. Details on sample collection and processing for these measurements are available at https://bats.bios.edu/data/. Values for all environmental variables are in Dataset S1. To explore depth-related shifts in biology and carbon export, we partitioned samples based on their position in photic vs. aphotic zones. The photic zone is well established and operationally defined at BATS as ∼1-140 m (to the 0.1% PAR), transitioning to the aphotic zone (160-1000 m) below 0.1% PAR (40, 65).

Bulk POC flux was estimated from monthly surface-tethered Particle Interceptor Traps (PITs) that were deployed at 150, 200, and 300 m and collected particles for 3 d (20). Triplicate trap tubes were fitted with acid-cleaned 0.8 µm polycarbonate filters at the bottom and filled with poisoned seawater brine. Trap filters were processed based on standard BATS protocol (67) to determine bulk POC flux using C/N analyses (20). Mean Syndiniales relative abundance in the photic zone (1-140 m) was correlated with bulk POC flux at 150 m via Spearman rank (rs) correlation. Similar correlations were made with other important eukaryotic groups in the photic zone, like Dinophyceae and Arthropoda. Though deployed at the same site, there was a ∼2-3 d lag between DNA collection and sediment trap recovery. Bulk POC flux data at 150 m is also included in Dataset S1.

### DNA extraction, PCR, and 18S rRNA metabarcoding analysis

DNA extraction from Sterivex filters has been described in (33). Primers were used to amplify the 18S V4 hypervariable region: 5-CCAGCA[GC]C[CT]GCGGTAATTCC-3 and 5- ACTTTCGTTCTTGAT[CT][AG]A-3 (68). Initial PCR runs consisted of a denaturation step at 98°C for 30 s, 10 cycles at 98°C for 10 s, 53°C for 30 s, and 72°C for 30 s, followed by 15 cycles at 98°C for 10 s, 48°C for 30 s, and 72°C for 30 s, and a final elongation at 72°C for 10 min (33). PCR reactions (25 µl) were run with 2 µl of target DNA, 1 µl of each primer, and 12.5 µl of KAPA HiFi HotStart ReadyMix (Kapa Biosystems; Wilmington, MA, USA). A second PCR was performed using dual Illumina indices as per the Illumina Nextera XT Index Kit. Amplicon sequencing was carried out on a MiSeq (2 x 250 bp) at the Center for Genome Research and Biocomputing at Oregon State University.

Primers were removed from demultiplexed reads using Cutadapt (69). Amplicon sequence variants (ASVs) were assigned to trimmed 18S reads using paired-end DADA2 in QIIME 2 (70). Taxonomy was assigned using a Naïve Bayes classifier trained with the Protistan Ribosomal Reference (or PR2) database (Version 5.0.1; (71)) and trimmed to the primer region (72). The current PR2 release included several updates, changing kingdom to domain and adding a new subdivision level for a total of 9 taxonomic levels (73). The PR2 database includes some metazoans assignments, which were considered here to explore potential Syndiniales-zooplankton relationships. Resulting taxonomy, ASV count, and metadata files were imported into R (Version 4.2.1) using qiime2R (https://github.com/jbisanz/qiime2R) and merged with phyloseq (74). Several groups were filtered from the dataset, including Craniata, Streptophyta, Rhodophyta, Bacteria, and Unassigned reads at the division level. Samples with less than 5,000 read counts and global singletons (ASVs present once) were also removed. Count tables were rarefied to the minimum read count (15,063 reads). The filtered ASV taxonomy and read counts table is provided in Dataset S3.

Changes in mean relative abundance with depth and season were observed using stacked bar plots in the R package microeco (75). Taxonomy was visualized at the class level, as well as order and family (clade) level within Syndiniales. Subsequently, phyloseq objects were trimmed to include only Syndiniales ASVs to focus on their population dynamics. Spatial and temporal trends in Syndiniales composition were observed via principial coordinates analysis (PCoA) using Bray-Curtis dissimilarity (76) and tested for significance with permutational multivariate analysis of variance (PERMANOVA) or one-way Analysis of Variance (ANOVA). Mean Shannon diversity index was estimated for samples in the photic vs. aphotic zones using the estimate_richness function in phyloseq (74).

### Partial least squares regression

The influence of environmental (predictor) variables on Syndiniales read counts (response variable) was determined with partial least squares regression (PLSR) using the R package mdatools (77). An initial PLSR model was generated using rarefied Syndiniales read counts. Predictor variables were centered and standardized. Outliers were detected using a Data Driven robust method (78) and removed (SI Appendix, Fig. 5). After running a final PLSR model, Root Mean Squared Error (RMSE) plots were used to select the optimal number of components (SI Appendix, Fig. 5). In this case, four components were optimal. Regression coefficients and variable influence on the projection (VIP) scores were estimated for each predictor variable (SI Appendix, Fig. 5). VIP scores > 1 are considered most important to the model (77). Spearman rank (rs) correlations were used to further explore predictor-response effects.

### Network analysis

Network analysis was applied to observe associations between Syndiniales and potential host organisms throughout the water column. Twelve separate networks were constructed, one for each depth, using the SParse InversE Covariance estimation for Ecological Associations and Statistical Inference (SPIEC-EASI) package in R (Version 1.1.0; (79)). SPIEC-EASI aims to minimize spurious edges from compositional data and infer direct associations between ASVs (79). To minimize dense networks, 18S datasets were subsampled to include only the top 150 most abundant ASVs at each depth. ASV count tables were centered log-ratio (clr) transformed and networks were run using the Meinshausen–Buhlmann’s neighborhood selection method and an optimal sparsity threshold of 0.05 (80). Depth networks were filtered to include only positive and negative edges between Syndiniales ASVs and other 18S groups. Networks were visualized in Cytoscape (81). Mean degree, or the number of edges connected to a given ASV (47), were estimated for Syndiniales and potential host ASVs to indicate the specificity of such connections.

To observe the overlap of Syndiniales ASVs between depths and seasons, we constructed UpSet plots using the R package ComplexUpset (82) that were based on presence-absence data. We further estimated the percent of Syndiniales ASVs that were present in the photic and aphotic zones, which may indicate vertical transport of parasites in the water column. Percent of transported parasites were compared to ASVs from other major taxonomic groups in the photic zone (Arthropoda and Dinophyceae), as well as all eukaryotic ASVs.

## Acknowledgments and funding sources

We thank the entire BATS personnel and the captains and crew of the RV Atlantic Explorer for their logistical support collecting and processing samples. The work was supported by NSF award OCE-1756105 (BATS program), Simons Foundation International’s BIOS-SCOPE program, and NSF award OCE-2019589 for the Center for Chemical Currencies of a Microbial Planet (C-CoMP). This is C-CoMP publication #028.

## Data availability

All code and files needed to reproduce results from this study are available at https://github.com/sra34/BATS-parasites. The GitHub repository also includes metadata, network files, and raw QIIME 2 files needed for the analysis. 18S rRNA sequences have been deposited in the Sequence Read Archive under BioProject PRJNA769790.

---

Figure S1. Relative abundance of the top 10 18S class level groups at each depth and season. Values represent the mean at each depth for samples collected in winter (n = 117), spring (n = 102), summer (n = 229), and fall (n = 94). Seasonal delineations correspond to the position of the mixed layer depth. The “Others” category reflects 18S groups that were <2% relative abundance at any depth.

Figure S2. Stacked bar plots of Syndiniales relative abundance at the order level at each depth. Values represent the mean at each depth; n = 43-53. The “Others” category reflects Syndiniales groups that were <2% relative abundance at any depth.

Figure S3. Principal coordinates analysis based on Bray-Curtis dissimilarity of only Syndiniales ASVs, filtered for four respective depths (1, 120, 600 and 1000 m; n = 44-53). Sample color and shape correspond to season and collection year. The proportion of variance explained by the first two axes is indicated on the plot.

Figure S4. Box plots displaying environmental variables with depth at BATS. Values represent the mean ± SD at each depth (n = 43-53). Filled circles reflect outliers, i.e., the observation is 1.5 times the interquartile range less than the first quartile or greater than the third quartile. Units for each factor are displayed. Temp = temperature; PO4 = phosphate; NO3 = nitrate; NO2 = nitrite; SiO4 = silicate; *p*CO2 = partial pressure of carbon dioxide; Alk = alkalinity; TN = total nitrogen; POC = particulate organic carbon; PON = particulate organic nitrogen; TOC = total organic carbon; Bact = bacteria cell density.

Figure S5. Results from partial least squares regression (PLSR) model selection of Syndiniales read counts. Top left panel shows robust outlier detection, with samples (blue points) plotted based on decomposition of X-Y distances. Outliers are points outside of the dotted line and were removed prior to running the final PLSR model. Top right panel shows the Root Mean Squared Error (RMSE) of measured (red) and predicted (blue) read counts. The RMSE plot shows that after four components, the model becomes overfitted. Bottom left panel shows the linear relationship (R^2^) of predicted vs. actual read counts, along with the optimal number of components for the model (ncomp = 4). Bottom right shows coefficient values for the 15 predictor variables considered in the PLSR model. Oxygen (µmol kg^-1^); TOC = total organic carbon (µmol kg^-1^); Bacteria = bacteria density (10^8^ cells kg^-1^); POC = particulate organic carbon (µg kg^-1^); SiO4 = silicate (µmol kg^-1^); PON = particulate organic nitrogen (µg kg^-1^); *p*CO2 = partial pressure of carbon dioxide (µatm); PO4 = phosphate (µmol kg^-1^); NO3 = nitrate (µmol kg^-^ 1); Salinity (psu); Temp = temperature (°C); Alkalinity (µmol kg^-1^); TN = total nitrogen (µmol kg^-1^); NO2 = nitrite (µmol kg^-1^).

Figure S6. Node degree from each depth network for 18S groups that may be potential hosts of Syndiniales (Arthropoda, Dinophyceae, Polycystinea, Acantharea, and RAD-A/B). In this case, degree represents the number of different Syndiniales ASVs connected to a given host ASV. Values represent the mean ± SD at each depth.

Figure S7. Number of network edges at each depth between Syndiniales ASVs and potential hosts at the genus level within each class level 18S group. For each class level group, only genera that were connected to Syndiniales >3 times at any given depth are shown, with the remaining grouped into an “Others” category (gray). Missing depths indicate a lack of observed network edges between Syndiniales and a specific group. Genus level assignments are based on the PR2 database, with ASVs assigned to lowest possible taxonomy.

Figure S8. UpSet plot to visualize the intersection of Syndiniales ASVs (from the networks) between seasons. The number of Syndiniales ASVs within each season are on the left side. Points and lines between points indicate the intersection, while bar plots on the top panel represent the number of shared Syndiniales ASVs for each intersection. Overlap is based on presence-absence data.

## References

1. C. B. Field, M. J. Behrenfeld, J. T. Randerson, P. Falkowski, Primary Production of the Biosphere: Integrating Terrestrial and Oceanic Components. Science 281, 237–240 (1998).

2. C. A. Durkin, B. A. S. Van Mooy, S. T. Dyhrman, K. O. Buesseler, Sinking phytoplankton associated with carbon flux in the Atlantic Ocean. Limnology and Oceanography 61, 1172– 1187 (2016).

3. D. A. Siegel, T. DeVries, I. Cetinić, K. M. Bisson, Quantifying the Ocean’s Biological Pump and Its Carbon Cycle Impacts on Global Scales. Annual Review of Marine Science 15, 329–356 (2023).

4. M. W. Lomas, et al., Increased ocean carbon export in the Sargasso Sea linked to climate variability is countered by its enhanced mesopelagic attenuation. Biogeosciences 7, 57–70 (2010).

5. T. DeVries, The Ocean Carbon Cycle. Annual Review of Environment and Resources 47, 317–341 (2022).

6. M. Nowicki, T. DeVries, D. A. Siegel, Quantifying the Carbon Export and Sequestration Pathways of the Ocean’s Biological Carbon Pump. Global Biogeochemical Cycles 36, e2021GB007083 (2022).

7. L. Guidi, et al., Plankton networks driving carbon export in the oligotrophic ocean. Nature 532, 465–470 (2016).

8. D. K. Steinberg, M. R. Landry, Zooplankton and the Ocean Carbon Cycle. Annual Review of Marine Science 9, 413–444 (2017).

9. D. A. Siegel, et al., Global assessment of ocean carbon export by combining satellite observations and food-web models. Global Biogeochemical Cycles 28, 181–196 (2014).

10. C. A. Durkin, et al., Tracing the path of carbon export in the ocean though DNA sequencing of individual sinking particles. ISME J 16, 1896–1906 (2022).

11. K. D. Lafferty, Interacting Parasites. Science 330, 187–188 (2010).

12. T. G. Jephcott, T. Sime-Ngando, F. H. Gleason, D. J. Macarthur, Host–parasite interactions in food webs: Diversity, stability, and coevolution. Food Webs 6, 1–8 (2016).

13. C. De Vargas, et al., Eukaryotic plankton diversity in the sunlit ocean. Science 348, 1261605 (2015).

14. I. Rizos, et al., Beyond the limits of the unassigned protist microbiome: inferring large- scale spatio-temporal patterns of Syndiniales marine parasites. ISME COMMUN. 3, 1–11 (2023).

15. L. Guillou, et al., Widespread occurrence and genetic diversity of marine parasitoids belonging to Syndiniales (Alveolata). Environmental Microbiology 10, 3349–3365 (2008).

16. L. J. Clarke, S. Bestley, A. Bissett, B. E. Deagle, A globally distributed Syndiniales parasite dominates the Southern Ocean micro-eukaryote community near the sea-ice edge. ISME Journal 13, 734–737 (2019).

17. G. A. Ollison, S. K. Hu, L. Y. Mesrop, E. F. DeLong, D. A. Caron, Come rain or shine: Depth not season shapes the active protistan community at station ALOHA in the North Pacific Subtropical Gyre. Deep Sea Research Part I: Oceanographic Research Papers 170, 103494 (2021).

18. D. Boeuf, et al., Biological composition and microbial dynamics of sinking particulate organic matter at abyssal depths in the oligotrophic open ocean. Proceedings of the National Academy of Sciences 116, 11824–11832 (2019).

19. C. M. Preston, C. A. Durkin, K. M. Yamahara, DNA metabarcoding reveals organisms contributing to particulate matter flux to abyssal depths in the North East Pacific ocean. Deep Sea Research Part II: Topical Studies in Oceanography 173, 104708 (2020).

20. B. N. Cruz, S. Brozak, S. Neuer, Microscopy and DNA-based characterization of sinking particles at the Bermuda Atlantic Time-series Study station point to zooplankton mediation of particle flux. Limnology and Oceanography 66, 3697–3713 (2021).

21. B. Valencia, et al., Microbial communities associated with sinking particles across an environmental gradient from coastal upwelling to the oligotrophic ocean. Deep Sea Research Part I: Oceanographic Research Papers 179, 103668 (2022).

22. A. Z. Worden, et al., Rethinking the marine carbon cycle: Factoring in the multifarious lifestyles of microbes. Science 347, 127594 (2015).

23. M. Torres-Beltrán, T. Sehein, M. G. Pachiadaki, S. J. Hallam, V. Edgcomb, Protistan parasites along oxygen gradients in a seasonally anoxic fjord: A network approach to assessing potential host-parasite interactions. Deep-Sea Research Part II: Topical Studies in Oceanography 156, 97–110 (2018).

24. D. J. S. Montagnes, A. Chambouvet, L. Guillou, A. Fenton, Responsibility of microzooplankton and parasite pressure for the demise of toxic dinoflagellate blooms. Aquatic Microbial Ecology 53, 211–225 (2008).

25. D. W. Coats, M. G. Park, Parasitism of photosynthetic dinoflagellates by three strains of Amoebophrya (Dinophyta): parasite survival, infectivity, generation time, and host specificity. Journal of Phycology 38, 520–528 (2002).

26. T. G. Jephcott, et al., Ecological impacts of parasitic chytrids, syndiniales and perkinsids on populations of marine photosynthetic dinoflagellates. Fungal Ecology 19, 47–58 (2016).

27. J. Decelle, et al., Intracellular development and impact of a marine eukaryotic parasite on its zombified microalgal host. ISME J 16, 2348–2359 (2022).

28. W. Yih, D. Wayne Coats, Infection of Gymnodinium sanguineum by the dinoflagellate Amoebophrya sp.: Effect of nutrient environment on parasite generation time, reproduction, and infectivity. Journal of Eukaryotic Microbiology 47, 504–510 (2000).

29. M. G. Park, S. Kim, E. Y. Shin, W. Yih, D. W. Coats, Parasitism of harmful dinoflagellates in Korean coastal waters. Harmful Algae 30, S62–S74 (2013).

30. R. Siano, et al., Distribution and host diversity of Amoebophryidae parasites across oligotrophic waters of the Mediterranean Sea. Biogeosciences 8, 267–278 (2011).

31. L. Berdjeb, A. Parada, D. M. Needham, J. A. Fuhrman, Short-term dynamics and interactions of marine protist communities during the spring-summer transition. ISME Journal 12, 1907–1917 (2018).

32. D. K. Steinberg, et al., Overview of the US JGOFS Bermuda Atlantic Time-series Study (BATS): a decade-scale look at ocean biology and biogeochemistry. Deep Sea Research Part II: Topical Studies in Oceanography 48, 1405–1447 (2001).

33. L. Blanco-Bercial, et al., The protist community traces seasonality and mesoscale hydrographic features in the oligotrophic Sargasso Sea. Frontiers in Marine Science 9 (2022).

34. P. López-García, F. Rodríguez-Valera, C. Pedrós-Alió, D. Moreira, Unexpected diversity of small eukaryotes in deep-sea Antarctic plankton. Nature 409, 603–607 (2001).

35. M. C. Pernice, et al., Large variability of bathypelagic microbial eukaryotic communities across the world’s oceans. ISME J 10, 945–958 (2016).

36. M. T. Duret, R. S. Lampitt, P. Lam, Eukaryotic influence on the oceanic biological carbon pump in the Scotia Sea as revealed by 18S rRNA gene sequencing of suspended and sinking particles. Limnology and Oceanography 65, S49–S70 (2020).

37. F. Not, J. del Campo, V. Balagué, C. de Vargas, R. Massana, New insights into the diversity of marine picoeukaryotes. PLoS ONE 4, 1–5 (2009).

38. L. Santoferrara, A. Qureshi, A. Sher, L. Blanco-Bercial, The photic-aphotic divide is a strong ecological and evolutionary force determining the distribution of ciliates (Alveolata, Ciliophora) in the ocean. The Journal of eukaryotic microbiology, e12976 (2023).

39. M. D. DuRand, R. J. Olson, S. W. Chisholm, Phytoplankton population dynamics at the Bermuda Atlantic Time-series station in the Sargasso Sea. Deep Sea Research Part II: Topical Studies in Oceanography 48, 1983–2003 (2001).

40. C. A. Carlson, et al., Seasonal dynamics of SAR11 populations in the euphotic and mesopelagic zones of the northwestern Sargasso Sea. ISME J 3, 283–295 (2009).

41. R. J. Parsons, M. Breitbart, M. W. Lomas, C. A. Carlson, Ocean time-series reveals recurring seasonal patterns of virioplankton dynamics in the northwestern Sargasso Sea. ISME J 6, 273–284 (2012).

42. M. W. Lomas, et al., Two decades and counting: 24-years of sustained open ocean biogeochemical measurements in the Sargasso Sea. Deep Sea Research Part II: Topical Studies in Oceanography 93, 16–32 (2013).

43. U. Christaki, D.-I. Skouroliakou, L. Jardillier, Interannual dynamics of putative parasites (Syndiniales Group II) in a coastal ecosystem. Environmental Microbiology **n/a**.

44. M. Nagarkar, B. Palenik, Diversity and putative interactions of parasitic alveolates belonging to Syndiniales at a coastal Pacific site. Environmental Microbiology Reports **n/a**.

45. S. K. Hu, et al., Protistan grazing impacts microbial communities and carbon cycling at deep-sea hydrothermal vents. Proceedings of the National Academy of Sciences 118, e2102674118 (2021).

46. G. Lima-Mendez, et al., Determinants of community structure in the grobal plankton interactome. Science 348, 1262073 (2015).

47. L. Röttjers, K. Faust, From hairballs to hypotheses–biological insights from microbial networks. FEMS Microbiology Reviews 42, 761–780 (2018).

48. M. F. M. Bjorbækmo, A. Evenstad, L. L. Røsæg, A. K. Krabberød, R. Logares, The planktonic protist interactome: where do we stand after a century of research? ISME Journal, 9–11 (2019).

49. F. G. Blanchet, K. Cazelles, D. Gravel, Co-occurrence is not evidence of ecological interactions. Ecology Letters 23, 1050–1063 (2020).

50. U. Christaki, et al., Parasitic Eukaryotes in a Meso-Eutrophic Coastal System with Marked Phaeocystis globosa Blooms. Frontiers in Marine Science 4 (2017).

51. S. R. Anderson, E. L. Harvey, Temporal Variability and Ecological Interactions of Parasitic Marine Syndiniales in Coastal Protist Communities. mSphere 5, 10.1128/msphere.00209-20 (2020).

52. M. Goberna, M. Verdú, Cautionary notes on the use of co-occurrence networks in soil ecology. Soil Biology and Biochemistry 166, 108534 (2022).

53. S. Kim, et al., Genetic diversity of parasitic dinoflagellates in the genus Amoebophrya and its relationship to parasite biology and biogeography. Journal of Eukaryotic Microbiology 55, 1–8 (2008).

54. A. Skovgaard, Dirty tricks in the plankton: Diversity and role of marine parasitic protists. Acta Protozoologica 53, 51–62 (2014).

55. S. Zamora-Terol, A. Novotny, M. Winder, Reconstructing marine plankton food web interactions using DNA metabarcoding. Molecular Ecology 29, 3380–3395 (2020).

56. F. Not, R. Gausling, F. Azam, J. F. Heidelberg, A. Z. Worden, Vertical distribution of picoeukaryotic diversity in the Sargasso Sea. Environmental Microbiology 9, 1233–1252 (2007).

57. J. Bråte, et al., Radiolaria Associated with Large Diversity of Marine Alveolates. Protist 163, 767–777 (2012).

58. J. K. Dolven, et al., Molecular Diversity of Alveolates Associated with Neritic North Atlantic Radiolarians. Protist 158, 65–76 (2007).

59. T. Biard, Diversity and ecology of Radiolaria in modern oceans. Environmental Microbiology 24, 2179–2200 (2022).

60. M. A. Moran, et al., The Ocean’s labile DOC supply chain. Limnology and Oceanography, 1–15 (2022).

61. L. Blanco-Bercial, Metabarcoding Analyses and Seasonality of the Zooplankton Community at BATS. Frontiers in Marine Science 7 (2020).

62. C. A. Durkin, M. L. Estapa, K. O. Buesseler, Observations of carbon export by small sinking particles in the upper mesopelagic. Marine Chemistry 175, 72–81 (2015).

63. A. Chambouvet, et al., Interplay Between the Parasite Amoebophrya sp. (Alveolata) and the Cyst Formation of the Red Tide Dinoflagellate Scrippsiella trochoidea. Protist 162, 637– 649 (2011).

64. S. R. Anderson, S. Menden-Deuer, Growth, Grazing, and Starvation Survival in Three Heterotrophic Dinoflagellate Species. J Eukaryot Microbiol 64, 213–225 (2017).

65. D. A. Hansell, C. A. Carlson, Biogeochemistry of total organic carbon and nitrogen in the Sargasso Sea: control by convective overturn. Deep Sea Research Part II: Topical Studies in Oceanography 48, 1649–1667 (2001).

66. K. Bisson, D. A. Siegel, T. DeVries, Diagnosing Mechanisms of Ocean Carbon Export in a Satellite-Based Food Web Model. Frontiers in Marine Science 7 (2020).

67. A. H. Knap, et al., BATS Methods Manual, Version 4 (1997) (June 18, 2023).

68. T. Stoeck, et al., Multiple marker parallel tag environmental DNA sequencing reveals a highly complex eukaryotic community in marine anoxic water. Molecular Ecology 19, 21– 31 (2010).

69. M. Martin, Cutadapt removes adapter sequences from high-throughput sequencing reads. EMBnet.journal 17, 10–12 (2011).

70. B. J. Callahan, P. J. Mcmurdie, S. P. Holmes, Exact sequence variants should replace operational taxonomic units in marker-gene data analysis. The ISME Journal 11, 2639– 2643 (2016).

71. L. Guillou, et al., The Protist Ribosomal Reference database (PR2): A catalog of unicellular eukaryote Small Sub-Unit rRNA sequences with curated taxonomy. Nucleic Acids Research 41 (2013).

72. N. A. Bokulich, et al., Optimizing taxonomic classification of marker-gene amplicon sequences with QIIME 2’s q2-feature-classifier plugin. Microbiome 6, 90 (2018).

73. F. Burki, A. J. Roger, M. W. Brown, A. G. B. Simpson, The New Tree of Eukaryotes. Trends in Ecology & Evolution 35, 43–55 (2020).

74. P. J. McMurdie, S. Holmes, Phyloseq: An R Package for Reproducible Interactive Analysis and Graphics of Microbiome Census Data. PLoS ONE 8 (2013).

75. C. Liu, Y. Cui, X. Li, M. Yao, microeco: an R package for data mining in microbial community ecology. FEMS Microbiology Ecology 97, fiaa255 (2021).

76. J. Oksanen, et al., vegan: Community Ecology Package. R package version 2.5-2. Cran R 1, 2 (2018).

77. S. Kucheryavskiy, mdatools – R package for chemometrics. Chemometrics and Intelligent Laboratory Systems 198, 103937 (2020).

78. O. Ye. Rodionova, A. L. Pomerantsev, Detection of Outliers in Projection-Based Modeling. Anal. Chem. 92, 2656–2664 (2020).

79. Z. D. Kurtz, et al., Sparse and Compositionally Robust Inference of Microbial Ecological Networks. PLOS Computational Biology 11, e1004226 (2015).

80. 80. H. Liu, K. Roeder, L. Wasserman, Stability Approach to Regularization Selection (StARS) for High Dimensional Graphical Models in Advances in Neural Information Processing Systems, (Curran Associates, Inc., 2010).

81. P. Shannon, et al., Cytoscape: A Software Environment for Integrated Models of Biomolecular Interaction Networks. Genome Research 13, 2498–2504 (2003).

82. M. Krassowski, M. Arts, C. Lagger, Max, krassowski/complex-upset: v1.3.5 (2022) https://doi.org/10.5281/zenodo.7314197 (June 18, 2023).

