## Supplementary Information for "Syndiniales parasites drive species networks and are a biomarker for carbon export in the oligotrophic ocean"

### Supplementary Figures

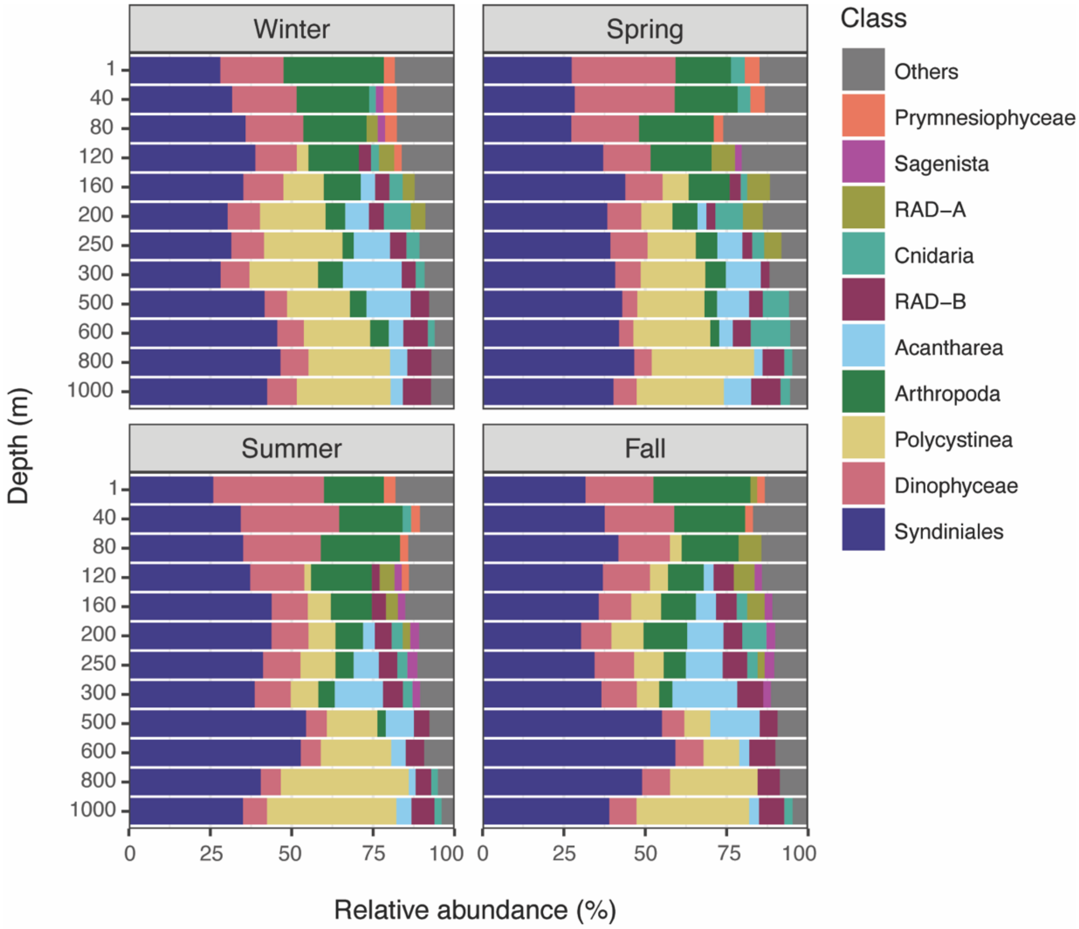


**Figure S1.** Relative abundance of the top 10 18S class level groups at each depth and season. Values represent the mean at each depth for samples collected in winter (n = 117), spring (n = 102), summer (n = 229), and fall (n = 94). Seasonal delineations correspond to the position of the mixed layer depth. The "Others" category reflects 18S groups that were <2% relative abundance at any depth.

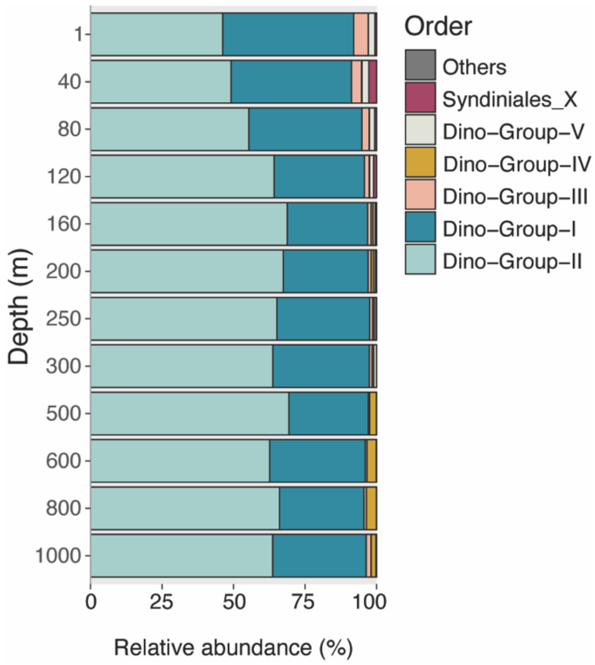


**Figure S2.** Stacked bar plots of Syndiniales relative abundance at the order level at each depth. Values represent the mean at each depth; n = 43-53. The “Others” category reflects Syndiniales groups that were <2% relative abundance at any depth.

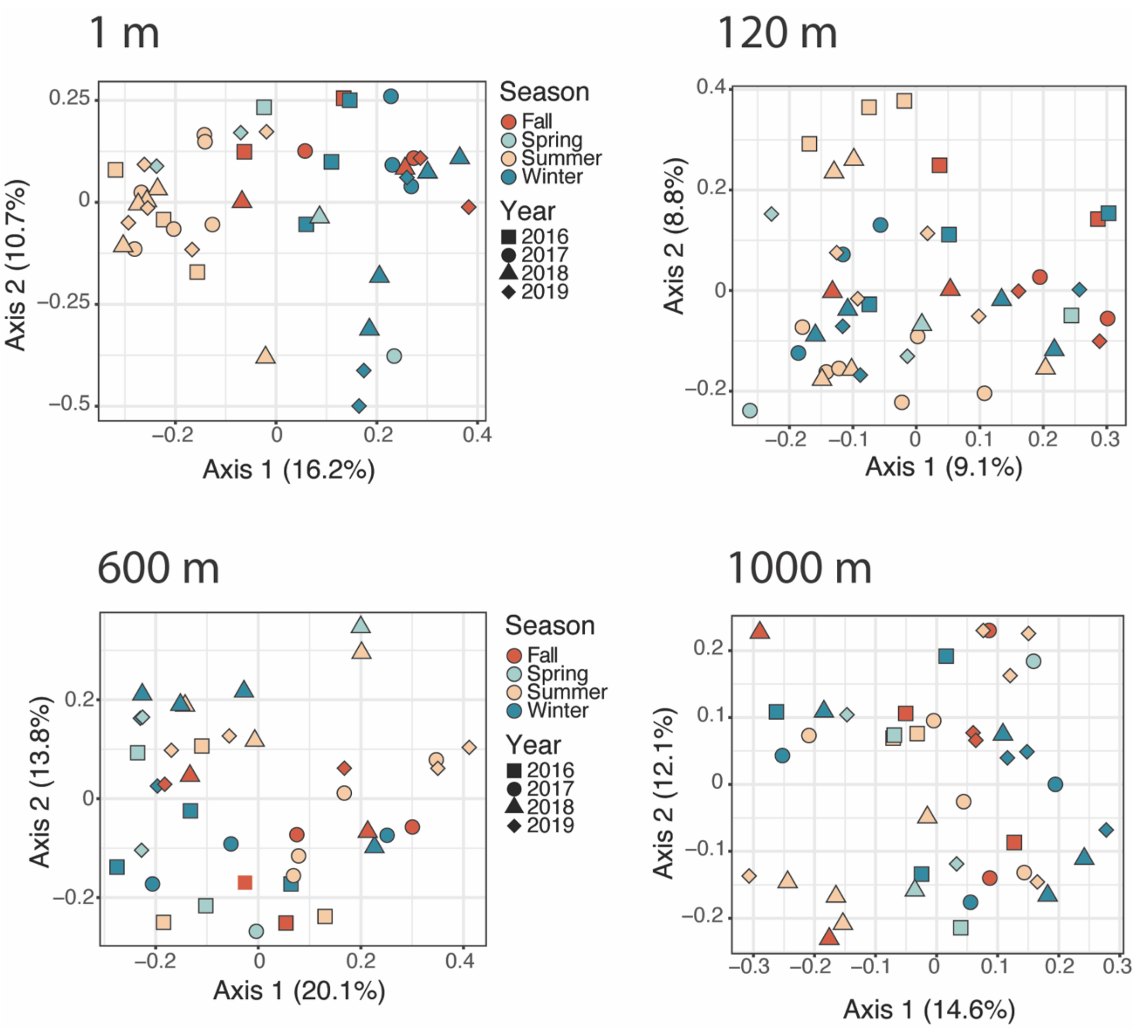


**Figure S3.** Principal coordinates analysis based on Bray-Curtis dissimilarity of only Syndiniales ASVs, filtered for four respective depths (1, 120, 600 and 1000 m; n = 44-53). Sample color and shape correspond to season and collection year. The proportion of variance explained by the first two axes is indicated on the plot.

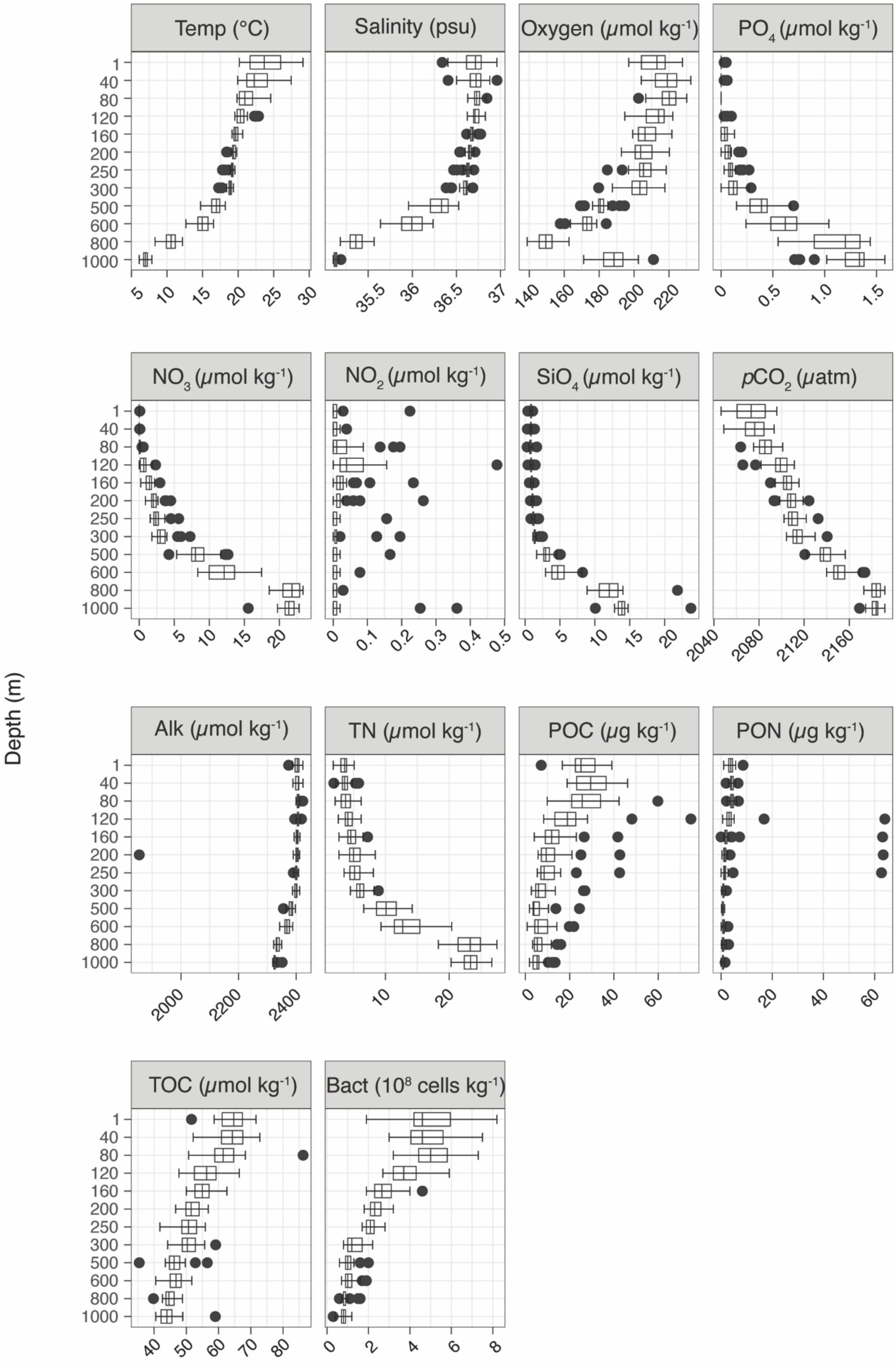
Temp (°C)
Salinity (psu)
Oxygen (μmol kg-1)
PO4 (μmol kg-1)
NO3 (μmol kg-1)
NO2 (μmol kg-1)
SiO4 (μmol kg-1)
pCO2 (μatm)
Alk (μmol kg-1)
TN (μmol kg-1)
POC (μg kg-1)
PON (μg kg-1)
TOC (μmol kg-1)
Bact (10^8 cells kg-1)
Depth (m)

**Figure S4.** Box plots displaying environmental variables with depth at BATS. Values represent the mean ± SD at each depth (n = 43-53). Filled circles reflect outliers, i.e., the observation is 1.5 times the interquartile range less than the first quartile or greater than the third quartile. Units for each factor are displayed. Temp = temperature; $PO_4$ = phosphate; $NO_3$ = nitrate; $NO_2$ = nitrite; $SiO_4$ = silicate; $pCO_2$ = partial pressure of carbon dioxide; Alk = alkalinity; TN = total nitrogen; POC = particulate organic carbon; PON = particulate organic nitrogen; TOC = total organic carbon; Bact = bacteria cell density.

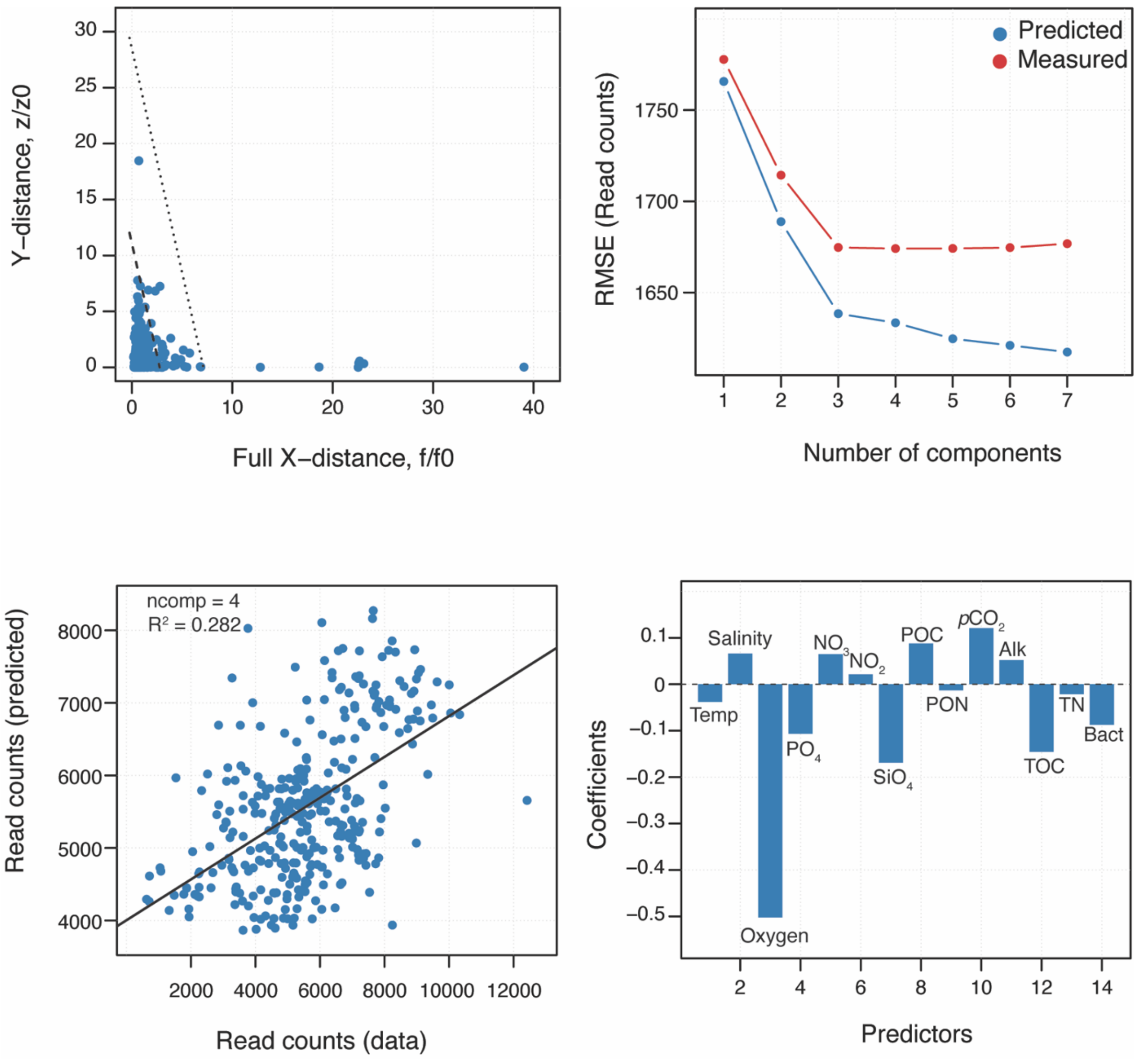


**Figure S5.** Results from partial least squares regression (PLSR) model selection of Syndiniales read counts. Top left panel shows robust outlier detection, with samples (blue points) plotted based on decomposition of X-Y distances. Outliers are points outside of the dotted line and were removed prior to running the final PLSR model. Top right panel shows the Root Mean Squared Error (RMSE) of measured (red) and predicted (blue) read counts. The RMSE plot shows that after four components, the model becomes overfitted. Bottom left panel shows the linear relationship ($R^2$) of predicted vs. actual read counts, along with the optimal number of components for the model (ncomp = 4). Bottom right shows coefficient values for the 15 predictor variables considered in the PLSR model. Oxygen (µmol kg$^{-1}$); TOC = total organic carbon (µmol kg$^{-1}$); Bacteria = bacteria density ($10^8$ cells kg$^{-1}$); POC = particulate organic carbon (µg kg$^{-1}$); $SiO_4$ = silicate (µmol kg$^{-1}$); PON = particulate organic nitrogen (µg kg$^{-1}$); $pCO_2$ = partial pressure of carbon dioxide (µatm); $PO_4$ = phosphate (µmol kg$^{-1}$); $NO_3$ = nitrate (µmol kg$^{-1}$); Salinity (psu); Temp = temperature (ºC); Alkalinity (µmol kg$^{-1}$); TN = total nitrogen (µmol kg$^{-1}$); $NO_2$ = nitrite (µmol kg$^{-1}$).

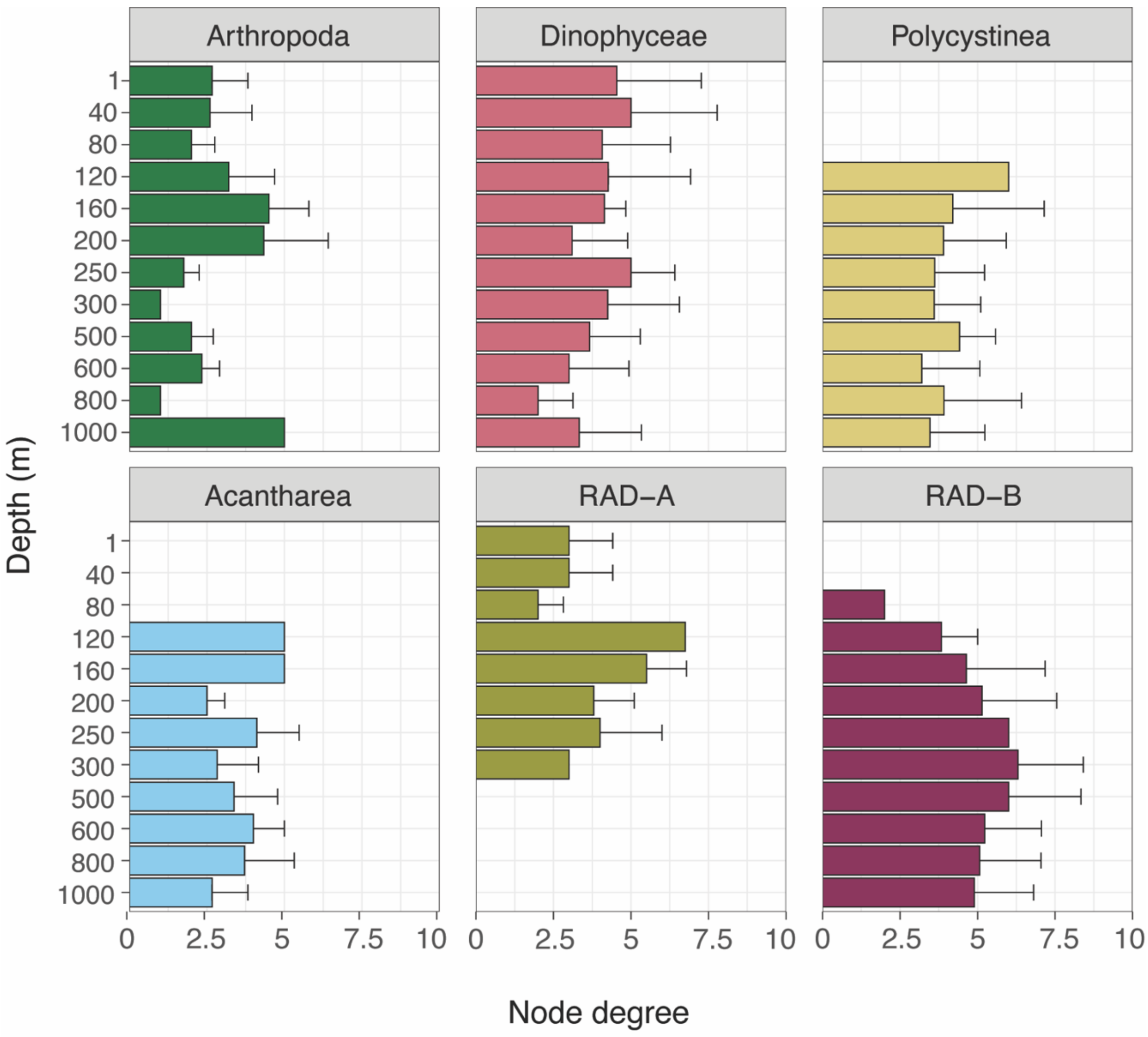


**Figure S6.** Node degree from each depth network for 18S groups that may be potential hosts of Syndiniales (Arthropoda, Dinophyceae, Polycystinea, Acantharea, and RAD-A/B). In this case, degree represents the number of different Syndiniales ASVs connected to a given host ASV. Values represent the mean ± SD at each depth.

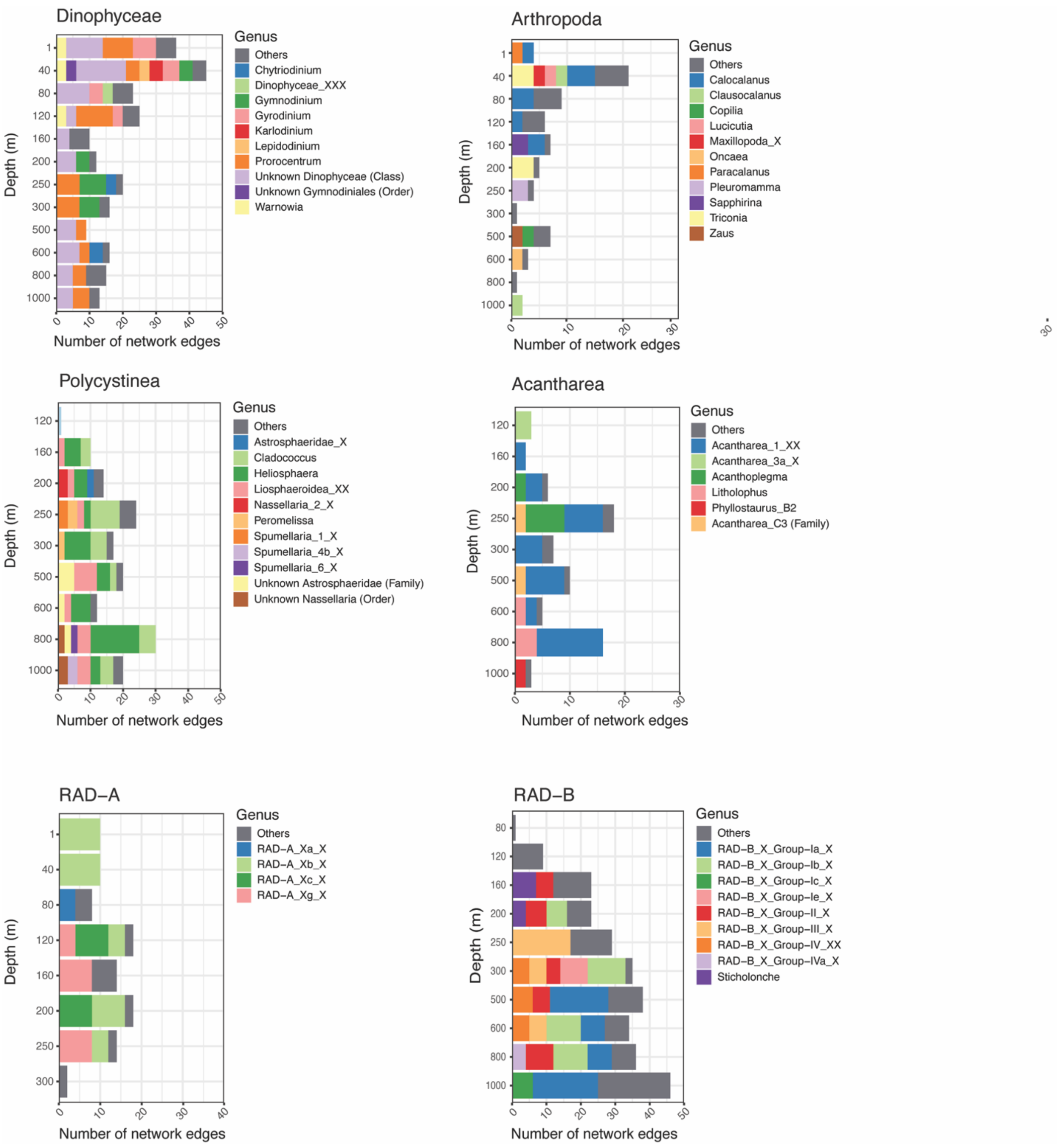


**Figure S7.** Number of network edges at each depth between Syndiniales ASVs and potential hosts at the genus level within each class level 18S group. For each class level group, only genera that were connected to Syndiniales >3 times at any given depth are shown, with the remaining grouped into an "Others" category (gray). Missing depths indicate a lack of observed network edges between Syndiniales and a specific group. Genus level assignments are based on the PR2 database, with ASVs assigned to lowest possible taxonomy.

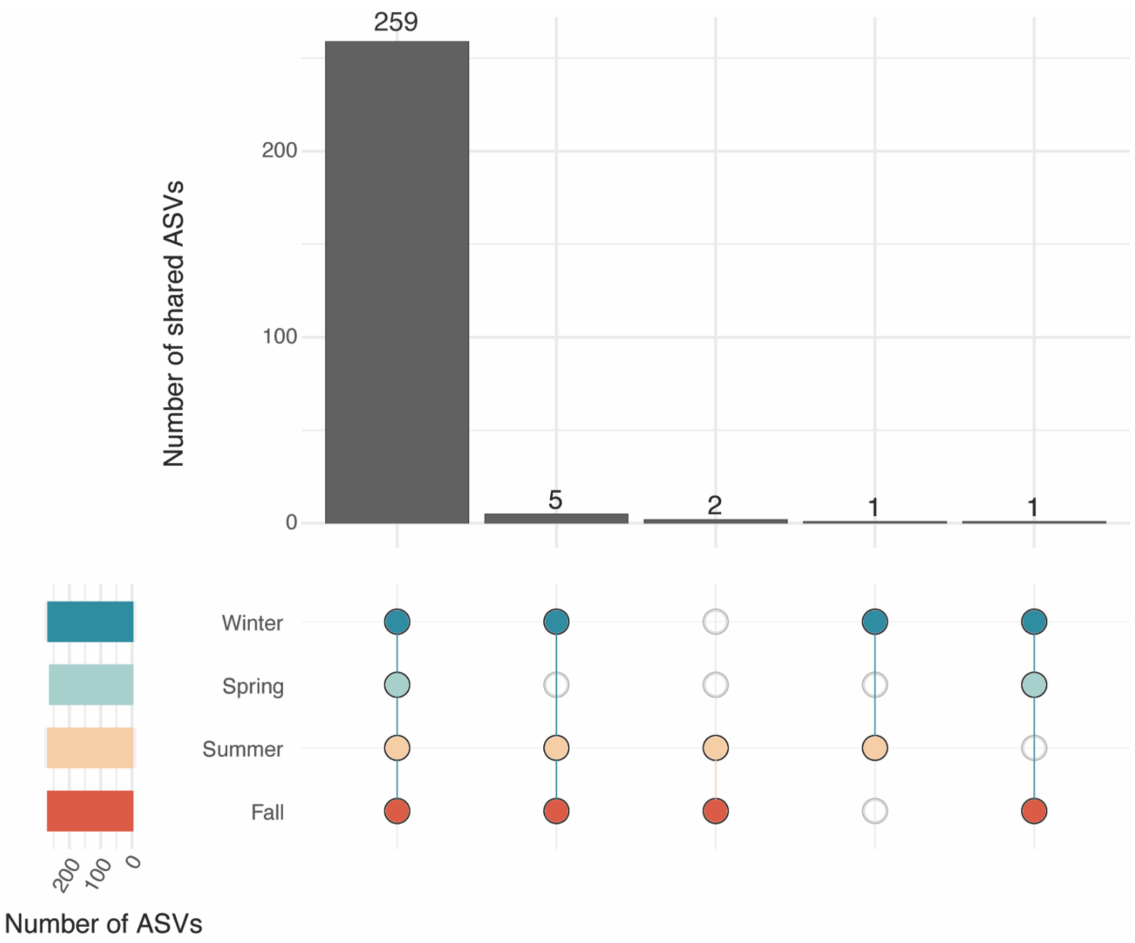


**Figure S8.** UpSet plot to visualize the intersection of Syndiniales ASVs (from the networks) between seasons. The number of Syndiniales ASVs within each season are on the left side. Points and lines between points indicate the intersection, while bar plots on the top panel represent the number of shared Syndiniales ASVs for each intersection. Overlap is based on presence-absence data.

**See Supplementary files for Datasets S1-S3**

**Dataset S1.** Environmental parameters measured each month at BATS. Samples are distinguished by nominal sampling depth (1-1000 m), vertical zone (aphotic vs. photic), and season. Mixed layer depth (MLD_dens125) was calculated as surfaced density + 0.125. Deep chlorophyll maximum (DCM) depth is also provided. Measurements of POC flux at 150 m (via sediment traps) are on a separate sheet. POC flux data were collected at the same site but different day (~2-3 d lag). Triplicate POC flux measurements were taken from a single trap (C1-C3). The mean and SD of triplicate measurements are shown (C_avg and C_sd). Sediment trap data was not available at 150 m for samples collected in March 2016, December 2017, July 2017, and March 2018. Season is also provided for each flux measurement.

**Dataset S2.** Summary of all significant ASV edges (or correlations) derived from SPIEC-EASI network analysis of twelve discrete depths. Edge information for each pairing (ASV1 and ASV2) includes taxonomy (domain, class-species), edge weight (strength of correlation), and sign (positive or negative). Network associations are also distinguished by depth. Depth networks filtered to only show significant edges between Syndiniales and potential host groups are on a separate sheet.

**Dataset S3.** Filtered taxonomic assignments and raw sequence counts for eukaryotic ASVs from monthly samples collected at BATS. Sample names are the same as in the metadata file. Taxonomy was assigned using the latest release from the Protistan Ribosomal Reference (PR2) database, which has 9 taxonomic levels (domain, supergroup, division, subdivision, class, order, family, genus, and species). Reads have been filtered to remove unwanted groups (see Materials and Methods) and singletons (ASVs observed only once). Reads counts were also rarefied to the minimum read count (15,063).
